## Supplementary Figure 1-8 for "mTORC1/S6K1 signaling promotes sustained oncogenic translation through modulating CRL3^IBTK^-mediated non-degradative ubiquitination of eIF4A1"

**This PDF file includes:**

Figures. S1 to S8

**Other Supplementary Materials for this manuscript include the following:**

Table S1 to S8

**Supplementary Table 1.** High-confidence IBTK-interacting protein identified with AP-MS method.

**Supplementary Table 2.** High-confidence IBTK-interacting protein identified with BioID method.

**Supplementary Table 3.** Ubiquitinated peptide sequences of eIF4A1.

**Supplementary Table 4.** Phosphorylated peptide sequences of IBTK by mTOR.

**Supplementary Table 5.** Phosphorylated peptide sequences of IBTK by S6K1.

**Supplementary Table 6.** Sequence information.

**Supplementary Table 7.** Antibody information.

**Supplementary Table 8.** Cell cultures, chemicals, and Kits.

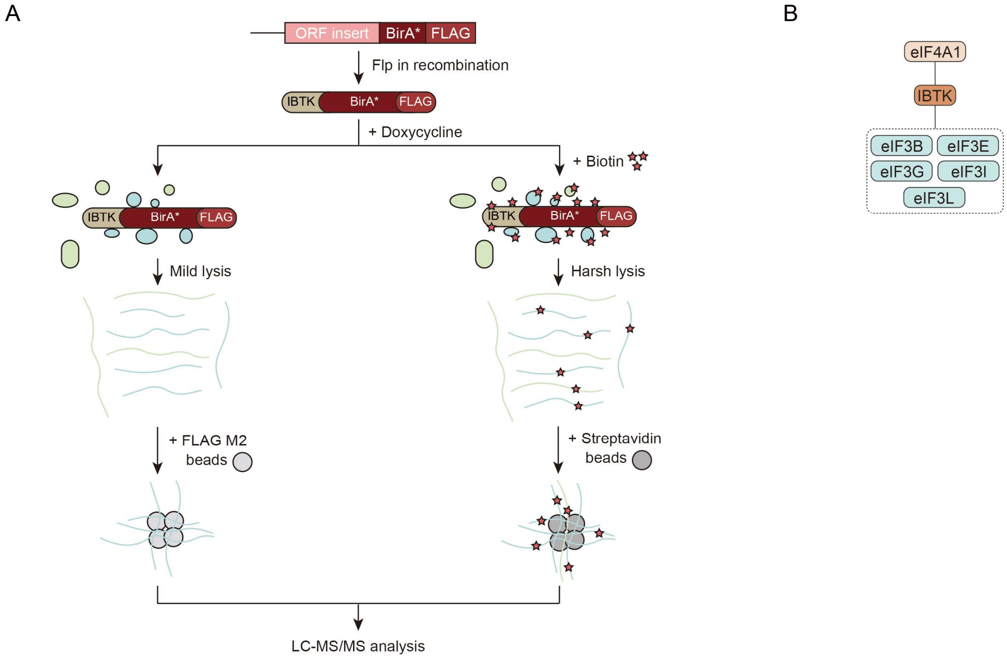


**Supplementary Figure 1. IBTK interacts with eIF4A1 in cells (related to Figure. 1).**

(**A**) A workflow for purification of IBTK protein complexes and identification potential IBTK interactors. The pcDNA5/FRT/TO vectors were modified to add a FLAG-tag and BirA* CDS. IBTK CDS was inserted to the MCS of this vector. The expression vector was then transfected into Flp-In T-REx 293 cells to establish inducible expressing isogenic cell lines. AP-MS and BioID methods were applied to purify IBTK protein complexes, respectively. The purified complexes were then subjected to mass-spectrometric sequencing.

(**B**) The eIF interactome of IBTK identified by BioID analysis. The full list of IBTK binding partners was showed in **Supplementary Table 2**.

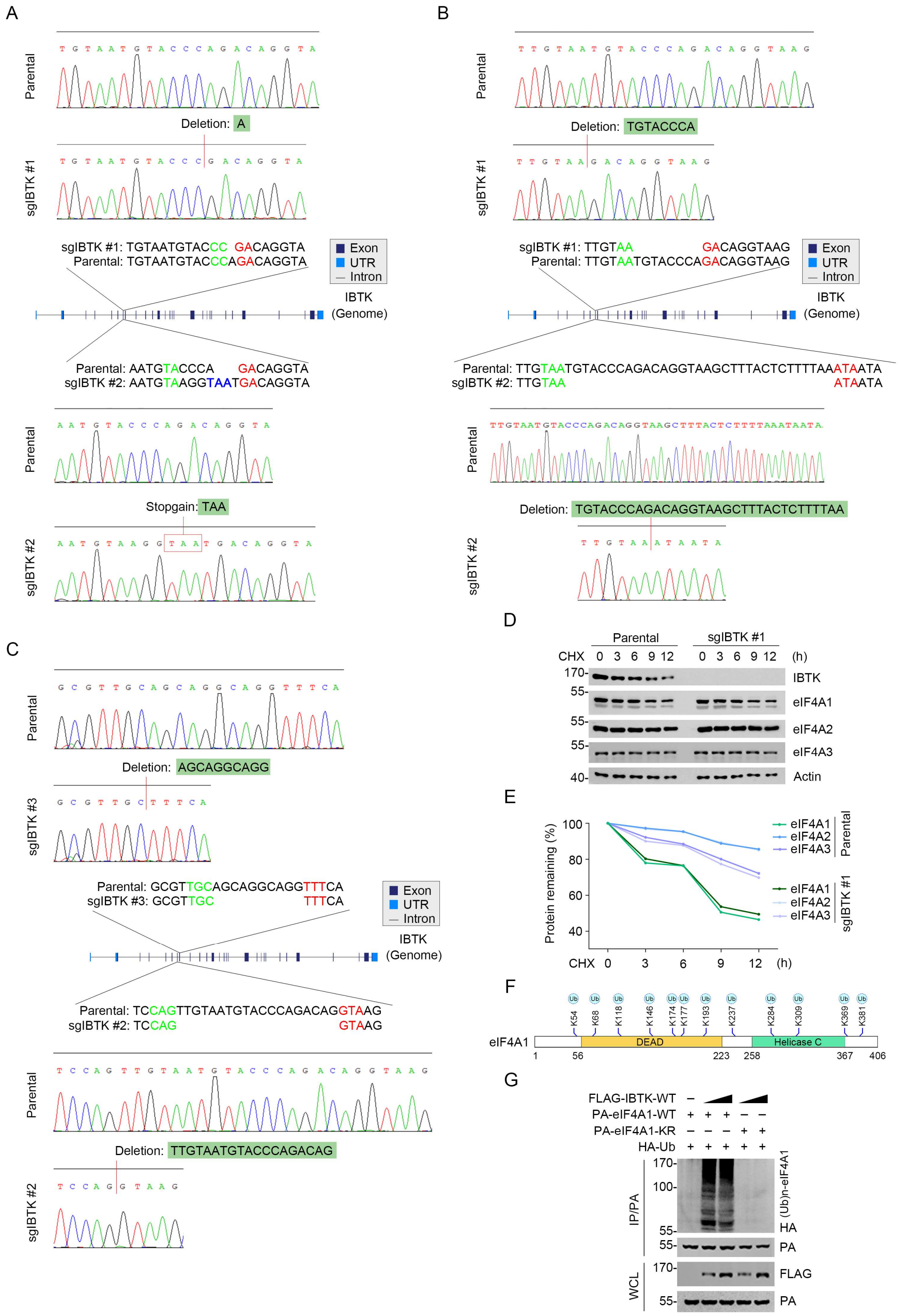


**Supplementary Figure 2. IBTK promotes non-degradative ubiquitination of eIF4A1 (related to Figure. 2).**

(**A**) Schematic of CRISPR/Cas9-mediated IBTK KO in SiHa cells and Sanger sequencing confirming that the IBTK gene was edited by sgRNA #1 or sgRNA #2 in SiHa cells.

(**B**) Schematic of CRISPR/Cas9-mediated IBTK KO in 293T cells and Sanger sequencing confirming that the IBTK gene was edited by sgRNA #1 or sgRNA #2 in 293T cells.

(**C**) Schematic of CRISPR/Cas9-mediated IBTK KO in H1299 cells and Sanger sequencing confirming that the IBTK gene was edited by sgRNA #1 or sgRNA #2 in H1299 cells.

(**D**, **E**) WB analysis of the indicated proteins in the WCLs from parental and IBTK-KO SiHa cells treated with cycloheximide (CHX,100 μg/ml) and harvested at different time points (**D**). At each time point, the intensity of eIF4A1, eIF4A2, and eIF4A3 was normalized to the intensity of Actin and then to the value at 0 h (**E**).

(**F**) Mass spectrometry analysis of the immunoprecipitated eIF4A1-Ub conjugate*s* reveals that eIF4A1 ubiquitination occurs at 12 lysine residues.

(**G**) WB analysis of the products of *in vivo* ubiquitination assays from 293T cells transfected with the indicated plasmids. Twelve ubiquitination sites in eIF4A1 were simultaneously mutated from lysine to arginine residues (eIF4A1-KR).

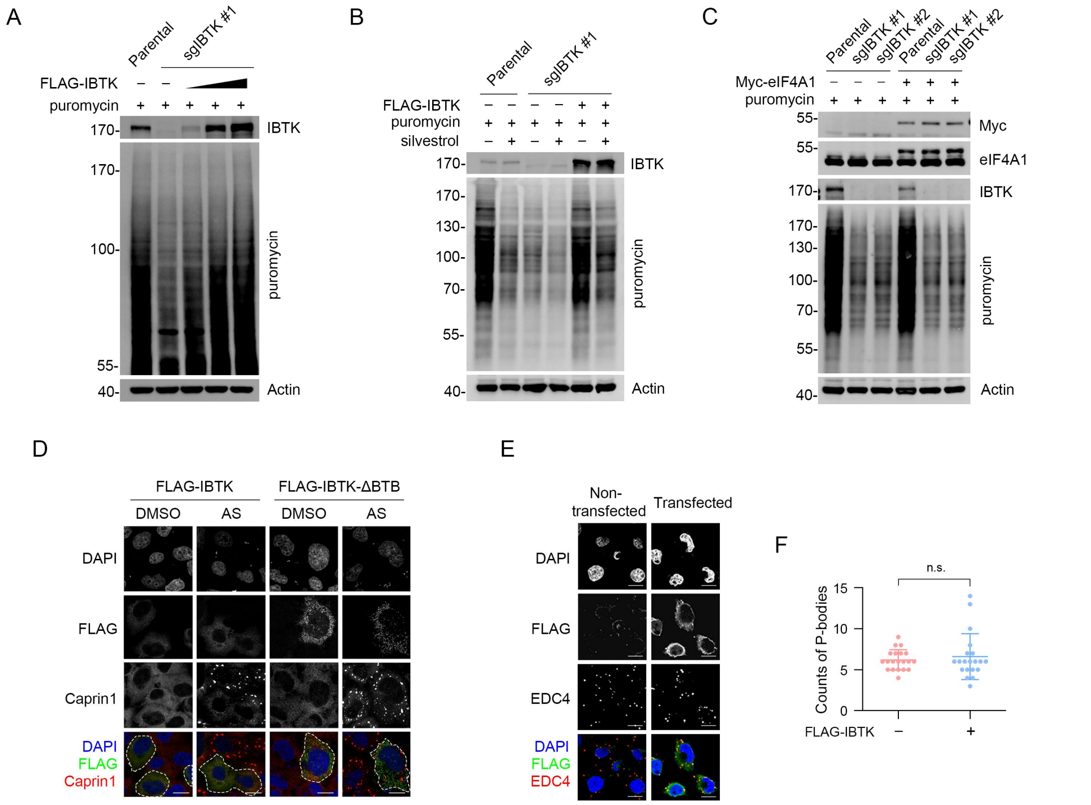


**Supplementary Figure 3. IBTK promotes nascent protein synthesis and cap-dependent translation initiation (related to Figure 3).**

(**A**) WB analysis of the indicated proteins in the WCLs from parental and IBTK-KO SiHa cells transfected with FLAG-IBTK and treated with puromycin (1.5 μM, 10 min) as indicated.

(**B**) WB analysis of the indicated proteins in the WCLs from parental and IBTK-KO SiHa cells transfected with FLAG-IBTK, treated with silvestrol (100 nM, 24 h) and puromycin (1.5 μM, 10 min).

(**C**) WB analysis of the indicated proteins in the WCLs from parental and IBTK-KO SiHa cells transfected with Myc-eIF4A1 and treated with puromycin (1.5 μM, 10 min).

(**D**) Representative IF images of SiHa cells transfected with the indicated plasmids, treated with DMSO or AS (100 μM, 2 h), and then stained with anti-FLAG (green), Caprin1 (red), and DAPI (blue). The cells successfully transfected with FLAG-tag plasmids are marked with white dashed lines. Scale bar, 20 μm.

(**E**) Representative IF images of HeLa cells transfected with FLAG-IBTK, stained with anti-FLAG (green), EDC4 (red), and DAPI (blue).

(**F**) P-bodies in (**E**) were quantified by counts of EDC4 puncta per cell (IBTK transfected vs non-transfected). Data are presented as means ± S.D. (n = 20).

*P* values are calculated using unpaired Student’s t-test in (**C**). n.s. non-significant.

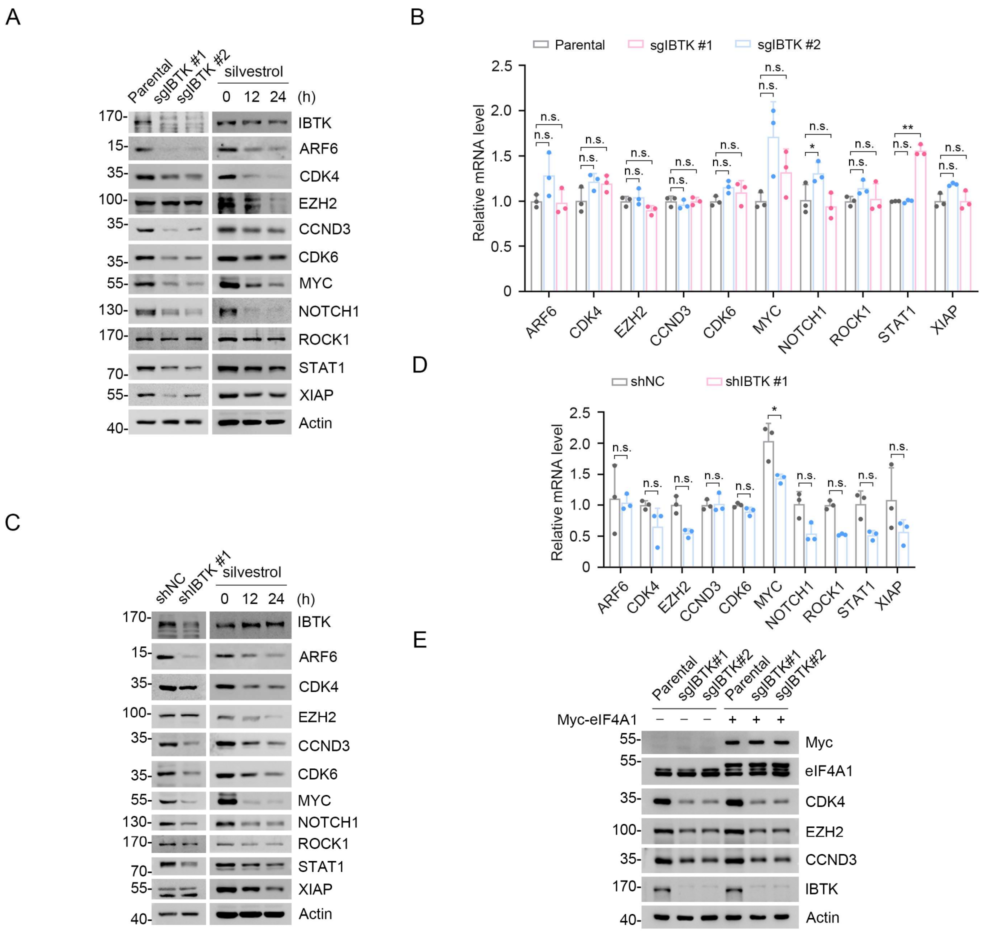


**Supplementary Figure 4. IBTK is indispensable for eIF4A1-related oncogene expression (related to Figure 4).**

(**A**) WB analysis of the indicated proteins in the WCLs from parental and IBTK-KO H1299 cells (left) and the WCLs from H1299 cells treated with silvestrol (100 nM) for the indicated times (right).

(**B**) RT-qPCR assessment of the mRNA expression of eIF4A1 targets in parental and IBTK-KO H1299 cells. The mRNA levels of *Actin* were used for normalization. Data are shown as means ± S.D. (n = 3).

(**C**) WB analysis of the indicated proteins in the WCLs from parental and IBTK-KD CT26 cells (left) and the WCLs from CT26 cells treated with silvestrol (100 nM) for the indicated times (right).

(**D**) RT-qPCR assessment of the mRNA expression of eIF4A1 targets in parental and IBTK-KD CT26 cells. The mRNA levels of *Actin* were used for normalization. Data are shown as means

± S.D. (n = 3).

(**E**) WB analysis of the indicated proteins in the WCLs from parental and IBTK-KO SiHa cells transfected with EV or Myc-eIF4A1.

*P* values are calculated using unpaired Two-way ANOVA test in (**B**, **D**). *p<0.05, **p<0.01, n.s. non-significant.

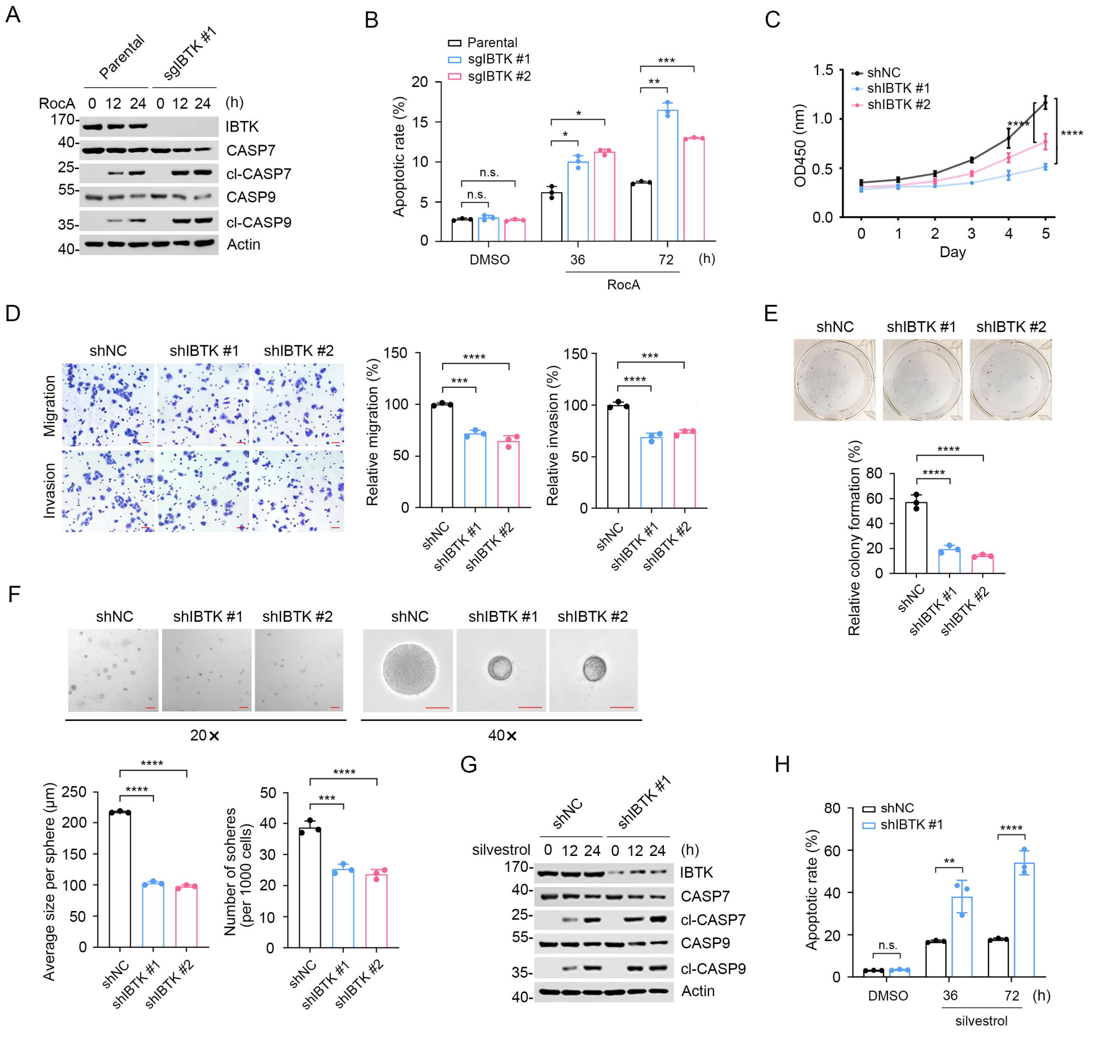


**Supplementary Figure 5. IBTK deficiency reduces neoplastic phenotypes in cancer cells (related to Figure 4).**

(**A**) WB analysis of the indicated proteins in the WCLs from parental and IBTK-KO SiHa cells treated with rocaglamide A (200 nM) for the indicated times.

(**B**) Parental and IBTK-KO SiHa cells were treated with rocaglamide A (200 nM) for the indicated times. Then, annexin-V-FITC/PI assays were used to stain the harvested cells, of which later flow cytometry analysis was performed. Data are shown as means ± S.D. (n=3).

(**C**) CCK-8 cell proliferation analysis of parental and IBTK-KD HeLa cells. Data are shown as means ± S.D. (n=3).

(**D**) Cell migration and invasion analysis of parental and IBTK-KD HeLa cells, and the quantitative analysis is shown on the right panel. Data are shown as means ± S.D. (n=3). Scale bar, 50 μm.

(**E**) Colony formation analysis of parental and IBTK-KD HeLa cells, and the quantitative data are shown as indicated. Data are shown as means ± S.D. (n=3).

(**F**) Sphere-formation analysis of parental and IBTK-KD HeLa cells. Representative pictures of HeLa cells after two weeks in three-dimensional culture are shown. Average size per sphere and number of spheres per 1000 cells were measured by ImageJ and result data are shown as means ± S.D. (n=3). Scale bar, 100 μm.

(**G**) WB analysis of the indicated proteins in the WCL from parental and IBTK-KD HeLa cells treated with silvestrol (100 nM) for the indicated times.

(**H**) Parental and IBTK-KD HeLa cells were treated with silvestrol (100 nM) for the indicated times. Then, annexin-V-FITC/PI assays were used to stain the harvested cells, of which later flow cytometry analysis was performed. Data are shown as means ± S.D. (n=3).

*P* values are calculated using One-way ANOVA test in (**D, E, F**), Two-way ANOVA test in (**B, C, H**). *p<0.05, **p<0.01, ***p<0.001, ****p<0.0001, n.s. non-significant.

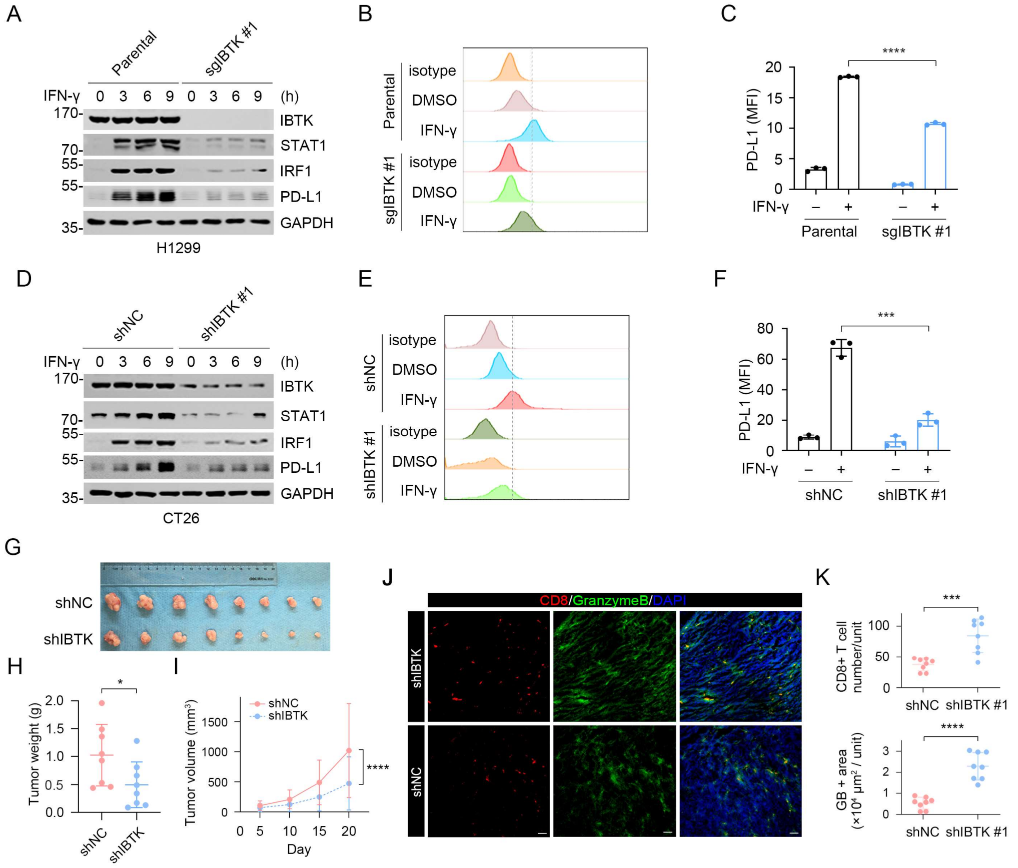


**Supplementary Figure 6. IBTK is required for IFN-γ-induced PD-L1 expression.**

(**A**) WB analysis of the indicated proteins in the WCLs from parental and IBTK-KO H1299 cells treated with IFN-γ (200 ng/ml) for the indicated times.

(**B**, **C**) PD-L1 mean fluorescence intensity (MFI) measurement of parental and IBTK-KO H1299 cells treated with DMSO or IFN-γ (200 ng/ml) for 24 h. Representative profiles (**B**) and MFI (**C**) are shown. Data are shown as means ± S.D. (n=3).

(**D**) WB analysis of the indicated proteins in the WCLs from parental and IBTK-KD CT26 cells treated with IFN-γ (200 ng/ml) for the indicated times.

(**E**, **F**) PD-L1 MFI measurement of parental or IBTK-KD CT26 cells treated with DMSO or IFN-γ (200 ng/ml) for 24 h. Representative profiles (**E**) and MFI (**F**) are shown. Data are shown as means ± S.D. (n=3).

(**G**-**I**) Parental and IBTK-KD CT26 cells were injected s.c. into the right flank of BALB/c mice. The tumors in each group at day 20 were harvested and photographed (**G**). The quantitative data of tumor weights (**H**) and tumor volumes (**I**) are shown. Data are shown as means ± S.D. (n=8).

(**J**, **K**) Immunostaining of CD8 and granzyme B in CT26 tumor mass. Scale bar, 200 μm. The number of $CD8^+$ T cells and the intensity of granzyme B were quantified using ImageJ and shown in (**K**). Data are shown as means ± S.D. (n=8).

*P* values are calculated using unpaired Student's t-test in (**H**, **K**), Two-way ANOVA test in (**C, I, F**). *p<0.05, ***p<0.001, ****p<0.0001, n.s. non-significant.

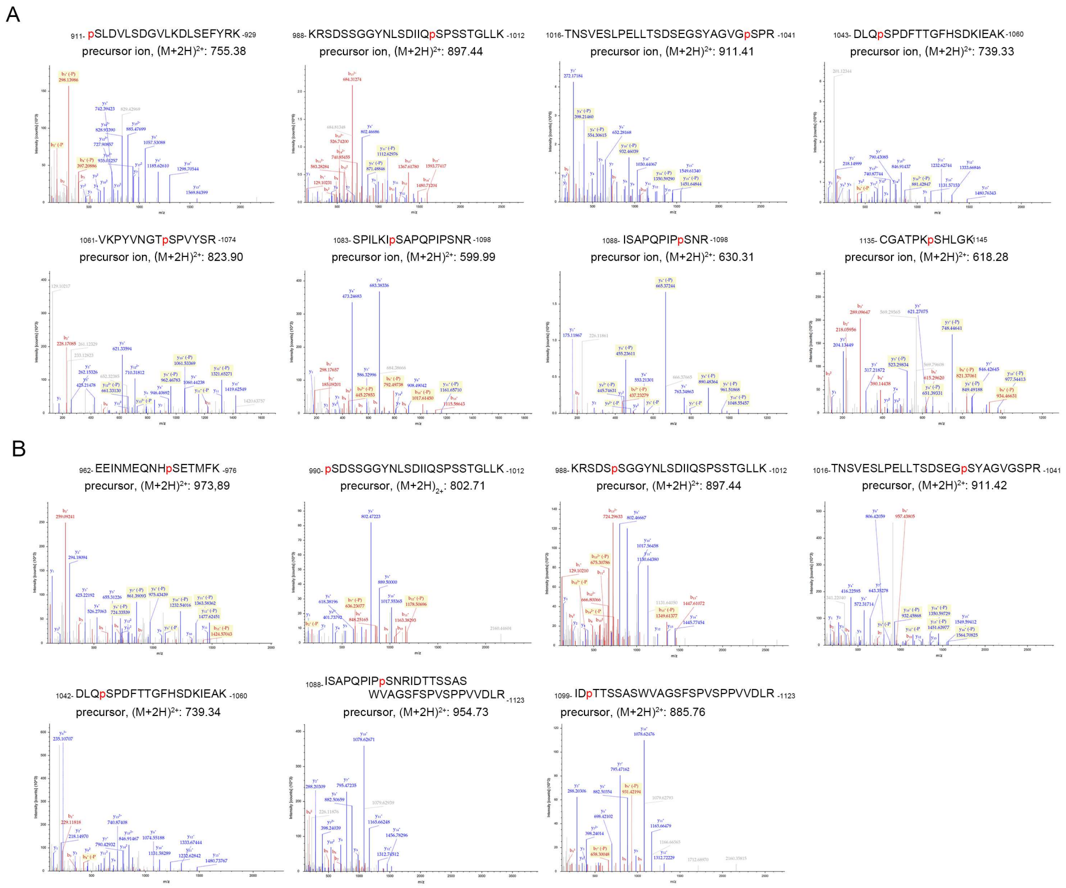


**Supplementary Figure 7. mTOR and S6K1 phosphorylates IBTK *in vitro* (related to Figure 5).**

(**A**) The MS spectra correspond to phosphorylated IBTK peptides in an *in vitro* mTOR/mLST8 phosphorylation assay.

(**B**) The MS spectra correspond to phosphorylated IBTK peptides in an *in vitro* S6K1 phosphorylation assay.

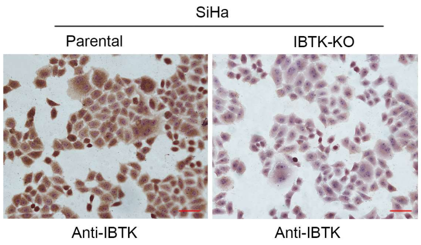


**Supplementary Figure 8. Representative images of IBTK IHC staining in parental and IBTK-KO SiHa cells (related to Figure 7).** Scale bar,100 μm.
